# Mouse suppressyn-like 1 suppresses membrane fusion through envelope glycoprotein recognition

**DOI:** 10.64898/2026.07.22.739964

**Authors:** Jun Sugimoto, Danny J Schust, So Nakagawa, Masateru Hiyoshi, Masumichi Saito, Makiko Sugimoto, Takeshi Nagamatsu, Haruko Takahashi, Yusuke Sotomaru, Yoshihiro Jinno, Yoshiki Kudo

## Abstract

Cell-cell fusion is essential for placental development and is mediated by endogenous retrovirus (ERV)-derived fusogens known as syncytins. However, how ERV-derived proteins negatively regulate membrane fusion remains largely unknown.

Here, we identify a previously uncharacterized murine ERV envelope-derived protein, mouse suppressyn-like 1 (mSUPYNL1), that suppresses syncytin-mediated membrane fusion through a mechanism distinct from that of placental human suppressyn (hSUPYN). Unlike hSUPYN, which acts through receptor interference, mSUPYNL1 inhibits both murine and human syncytin-mediated fusion independently of receptor usage by associating with the surface (SU) subunits of multiple syncytin envelope glycoproteins, revealing a receptor-independent mechanism of fusion suppression.

This mechanism extends beyond endogenous fusogens. mSUPYNL1 also associates with the SU glycoprotein (gp46) of Human T-cell Leukemia Virus type 1 (HTLV-1) and suppresses Env-dependent syncytium formation, whereas hSUPYN showed no detectable antiviral activity in this assay. These findings identify mSUPYNL1 as a broad-spectrum inhibitor of envelope glycoprotein-mediated membrane fusion.

Analysis of mSUPYNL1 knockout mice revealed that, in contrast to the placenta-restricted expression of hSUPYN, mSUPYNL1 was broadly expressed, with its most prominent localization in decidual stromal and vascular endothelial cells of the pregnant uterus, as well as in hematopoietic tissues such as the spleen and thymus.

Together, our findings uncover an evolutionarily distinct class of ERV-derived fusion suppressors that function through envelope glycoprotein recognition instead of receptor interference. Our study expands current models of ERV domestication by demonstrating that retroviral envelope proteins have been independently co-opted not only to promote membrane fusion but also to restrain it, thereby linking placental biology, antiviral defense, and host evolution.

**HIGHLIGHTS:**

- mSUPYNL1 is an endogenous retrovirus-derived membrane fusion inhibitor
- mSUPYNL1 binds the SU domains of murine and human syncytins
- mSUPYNL1 suppresses HTLV-1 Env-mediated syncytium formation
- Direct envelope recognition enables receptor-independent fusion inhibition

## Introduction

Endogenous retroviruses (ERVs) are remnants of ancient retroviral infections that have become permanently integrated into vertebrate genomes ^1–3^. Although most ERVs have lost their ancestral viral functions, some have been co-opted to perform essential physiological roles in the host. Among these, ERV-derived envelope (Env) proteins represent striking examples of molecular domestication, having been repeatedly repurposed during mammalian evolution.

The best-characterized ERV-derived Env proteins are the syncytins^4–9^, which mediate trophoblast fusion during placental development. Despite their independent evolutionary origins, primates and rodents have each acquired distinct syncytins that perform nearly equivalent roles in placental morphogenesis^10–14^. Genetic and functional studies have established that syncytins are indispensable for trophoblast fusion and placental development^15–19^.

In contrast, the mechanisms that restrain trophoblast fusion remained poorly understood. We previously identified human suppressyn (hSUPYN) as the first ERV-derived inhibitor of cell fusion^20^. hSUPYN inhibits syncytin-1-mediated fusion through receptor interference by binding the shared receptor, ASCT2, and is expressed predominantly in unfused trophoblast populations, suggesting a role in restricting trophoblast fusion. These findings established receptor interference as the first endogenous mechanism by which an ERV-derived protein suppresses membrane fusion. Whether receptor-independent mechanisms of ERV-mediated fusion suppression exist, and whether such mechanisms have evolved outside primates, remained unknown.

Here, we identify a previously uncharacterized mouse ERV-derived envelope protein, mouse suppressyn-like 1 (mSUPYNL1) (accession number: PZ682522). Unlike hSUPYN, mSUPYNL1 suppresses membrane fusion independently of receptor usage by associating with the surface (SU) subunits of multiple syncytin envelope glycoproteins. This mechanism extends beyond endogenous fusogens, as mSUPYNL1 also associates with the HTLV-1 envelope glycoprotein and suppresses Env-dependent syncytium formation. Analysis of knockout mice further revealed an expression pattern distinct from that of hSUPYN, indicating evolutionary diversification of ERV-derived fusion suppressors.

Together, our findings identify a previously unrecognized mechanism of ERV-derived fusion suppression, demonstrating that retroviral envelope proteins have been independently co-opted to regulate membrane fusion through distinct molecular strategies.

## Results

### Identification of mSUPYNL1 as a broad-spectrum ERV-derived membrane fusion inhibitor

To identify ERV-derived proteins that negatively regulate membrane fusion, we systematically screened mouse placental RNA-seq datasets using the Genome-based Endogenous Viral Elements (gEVE) database^21^. Candidate envelope (Env) genes were prioritized based on placental expression and the presence of intact open reading frames. Fifteen candidate loci were identified, of which four (C7, C11, C14, and C15) were confirmed to be expressed in the placenta by RT-PCR and selected for functional characterization (Fig. 1a and Supplementary Fig. 1a).

**Figure 1.**
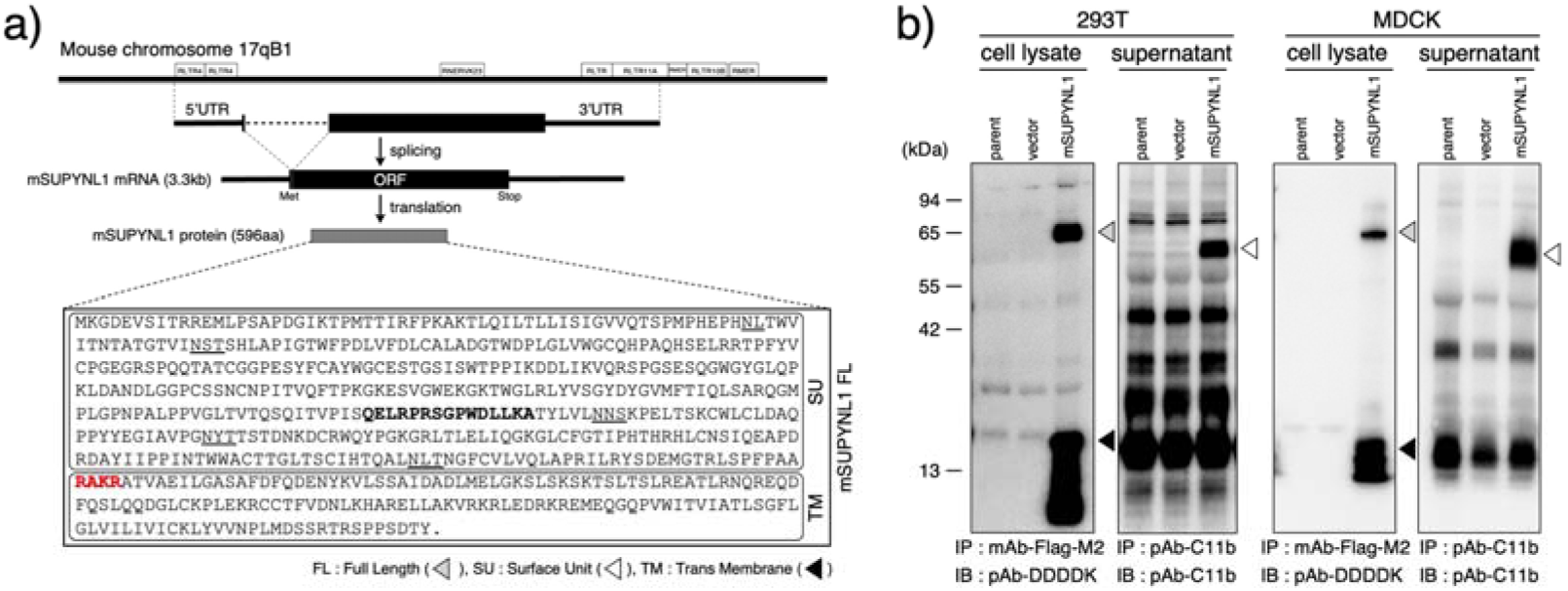
Genomic organization and maturation of the ERV-derived membrane fusion inhibitor mSUPYNL1. a, Schematic representation of the mSUPYNL1 locus on mouse chromosome 17qB1, together with its transcript and predicted protein structure. The full-length transcript encodes a 596-amino acid ERV-derived envelope glycoprotein containing a putative furin cleavage motif (RAKR)^36^, predicted to generate the surface (SU) and transmembrane (TM) subunits. Predicted N-linked glycosylation sites are underlined. The peptide sequence used for generation of the anti-mSUPYNL1 polyclonal antibody (pAb-C11b) is shown in bold. b, Expression and processing of mSUPYNL1. FLAG-tagged mSUPYNL1 was transiently expressed in 293T and MDCK cells and analyzed by immunoblotting. Cell lysates contained the full-length protein together with the TM subunit. Secreted SU protein was detected in conditioned medium following immunoprecipitation with the anti-mSUPYNL1 antibody (pAb-C11b), indicating proteolytic processing and extracellular release of the SU domain. Uncropped immunoblots are provided in Supplementary Fig. 6.

The four candidates exhibited distinct expression profiles. Whereas C15 showed placenta-restricted expression similar to syncytin-A and syncytin-B, C7, C11, and C14 were broadly expressed in multiple tissues while remaining detectable in the placenta. All four ORFs produced proteins of the expected molecular size after transient expression in cultured cells and displayed distinct intracellular localization patterns (Supplementary Fig. 1b, c). Functional screening revealed that only one candidate, C11, consistently inhibited syncytin-mediated membrane fusion. C11 almost completely suppressed multinucleated syncytium formation induced by both murine syncytin-A and syncytin-B, whereas the remaining candidates exhibited no detectable inhibitory activity. Unexpectedly, C11 also inhibited membrane fusion mediated by human syncytin-1 and syncytin-2, indicating that its anti-fusogenic activity is not species restricted (Supplementary Fig. 2).

Collectively, these findings identified C11 as a previously unrecognized ERV-derived membrane fusion inhibitor with broad activity against evolutionarily distinct syncytins. Based on these unique functional properties, we designated this protein mouse suppressyn-like 1 (mSUPYNL1).

### mSUPYNL1 suppresses membrane fusion through association with syncytin envelope glycoproteins

Genomic analysis revealed that mSUPYNL1 is an endogenous retroviral envelope gene located on mouse chromosome 17 and encodes a predicted 596-amino acid envelope glycoprotein (Fig. 1a). Sequence analysis showed the presence of a characteristic long terminal repeat (LTR) upstream of the coding region and identified feline leukemia virus Env as its closest homolog, consistent with its retroviral origin (Supplementary Fig. 3).

Biochemical analyses demonstrated that mSUPYNL1 undergoes canonical retroviral Env processing. Immunoblotting detected precursor and processed protein species, whereas immunoprecipitation of conditioned medium identified a secreted SU subunit, indicating proteolytic cleavage into surface (SU) and transmembrane (TM) components (Fig. 1b).

mSUPYNL1 potently suppressed membrane fusion mediated by both murine and human syncytins. Co-expression of mSUPYNL1 almost completely abolished syncytium formation induced by mouse syncytin-A and syncytin-B (Fig. 2a,b). Remarkably, equivalent inhibition was observed for human syncytin-1 and syncytin-2 (Supplementary Fig. 4a,b), demonstrating that the anti-fusogenic activity of mSUPYNL1 is conserved across evolutionarily distinct syncytins despite their different receptor specificities.

**Figure 2.**
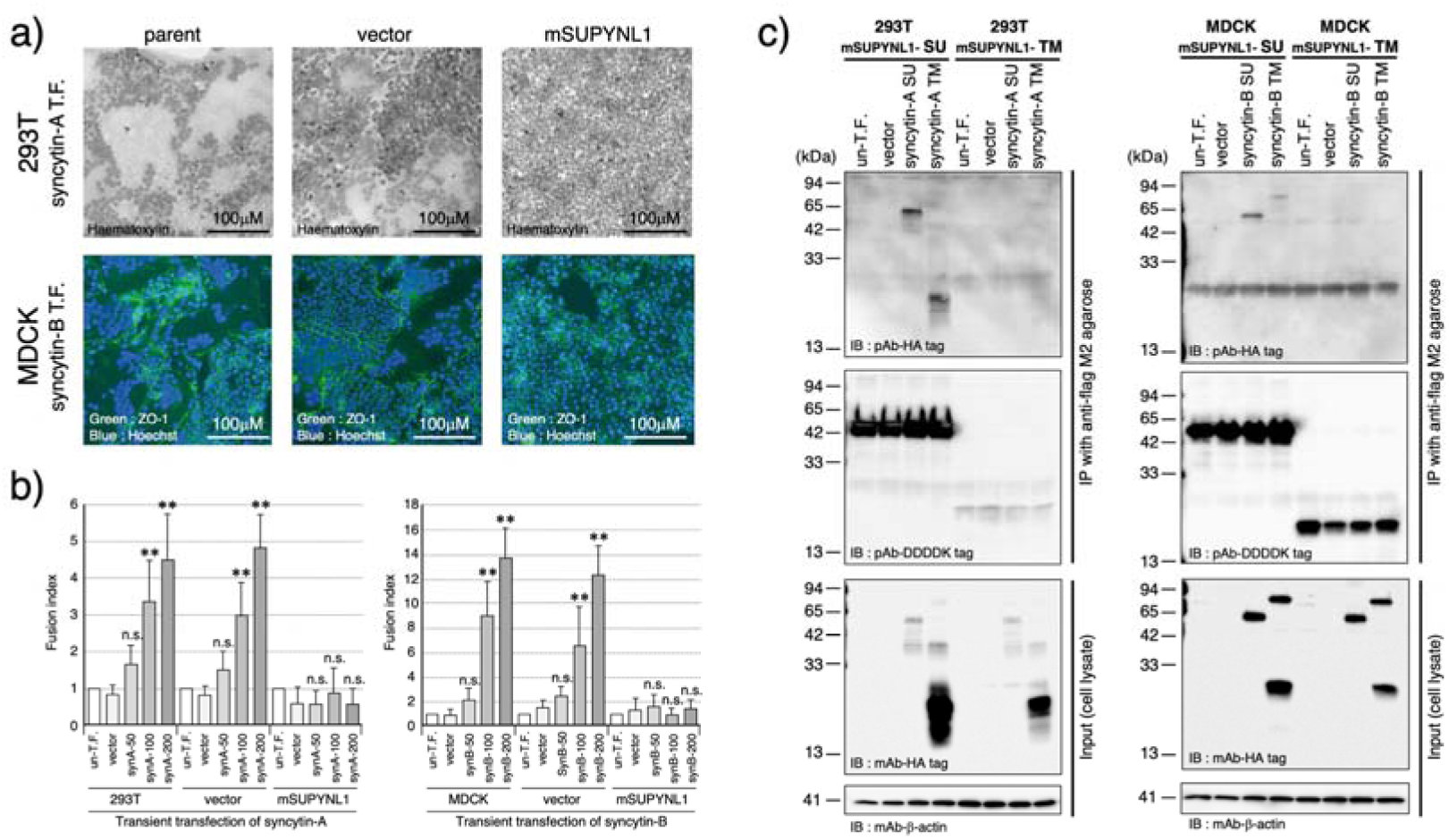
mSUPYNL1 suppresses syncytin-mediated membrane fusion through direct association with syncytin SU glycoproteins. a, Representative images of syncytin-mediated membrane fusion assays. Syncytin-A-induced fusion was examined in 293T cells and visualized by hematoxylin staining. Syncytin-B-induced fusion was analyzed in MDCK cells by immunofluorescence using anti-ZO-1 antibody to visualize cell boundaries. Forced expression of mSUPYNL1 markedly suppressed syncytium formation induced by both syncytin-A and syncytin-B. b, Quantification of membrane fusion by flow cytometry. Increasing amounts (50–200 ng) of syncytin expression plasmids were transfected, and fusion indices were calculated relative to untransfected cells. Data were analyzed by two-tailed non-repeated measures ANOVA. **, p < 0.01; n.s., not significant. c, mSUPYNL1 directly associates with syncytin SU glycoproteins. FLAG-tagged SU or TM domains of mSUPYNL1 were stably expressed in 293T or MDCK cells, followed by transient expression of HA-tagged SU or TM domains of syncytin-A or syncytin-B. Protein complexes were isolated by FLAG immunoprecipitation and analyzed by immunoblotting with anti-HA antibody. Input lysates demonstrate comparable protein expression, whereas immunoprecipitated fractions identify proteins directly associated with mSUPYNL1. Anti-FLAG and β-actin immunoblots served as immunoprecipitation and loading controls, respectively. Uncropped immunoblots are shown in Supplementary Fig. 7.

Because the syncytins examined utilize distinct cellular receptors, these findings suggested that fusion inhibition by mSUPYNL1 is unlikely to be explained by receptor competition. We therefore asked whether mSUPYNL1 instead associates with syncytin envelope glycoproteins.

Co-immunoprecipitation analyses revealed that the SU subunit of mSUPYNL1 preferentially associated with the SU domains of syncytin-A and syncytin-B, whereas little or no interaction was detected with their TM domains (Fig. 2c). A similar binding pattern was observed for human syncytin-1 and syncytin-2, indicating that mSUPYNL1 recognizes the SU domains of multiple retroviral envelope glycoproteins despite their independent evolutionary origins (Supplementary Fig. 4c,d).

Together, these findings demonstrate that mSUPYNL1 suppresses membrane fusion through association with retroviral envelope glycoproteins rather than receptor competition. This receptor-independent mechanism distinguishes mSUPYNL1 from hSUPYN and identifies envelope glycoprotein recognition as a previously unrecognized mechanism of ERV-derived membrane fusion suppression.

### mSUPYNL1 recognizes exogenous retroviral envelope glycoproteins and suppresses HTLV-1 Env-mediated membrane fusion

Because mSUPYNL1 associated with the envelope glycoproteins of multiple evolutionarily distinct syncytins, we asked whether this mechanism extends beyond endogenous retroviral fusogens. We therefore examined the human retrovirus, HTLV-1, whose cell-to-cell transmission depends on Env-mediated syncytium formation. mSUPYNL1 potently suppressed HTLV-1 Env-mediated membrane fusion. Extensive multinucleated syncytia developed in control cells and in cells expressing hSUPYN, whereas syncytium formation was almost completely abolished in mSUPYNL1-expressing cells (Fig. 3a,b). Quantitative analysis revealed an approximately 90% reduction in HTLV-1 Env-dependent membrane fusion relative to control cells (Fig. 3c). Notably, hSUPYN exhibited no detectable inhibitory activity, consistent with its receptor-dependent mechanism of action.

**Figure 3.**
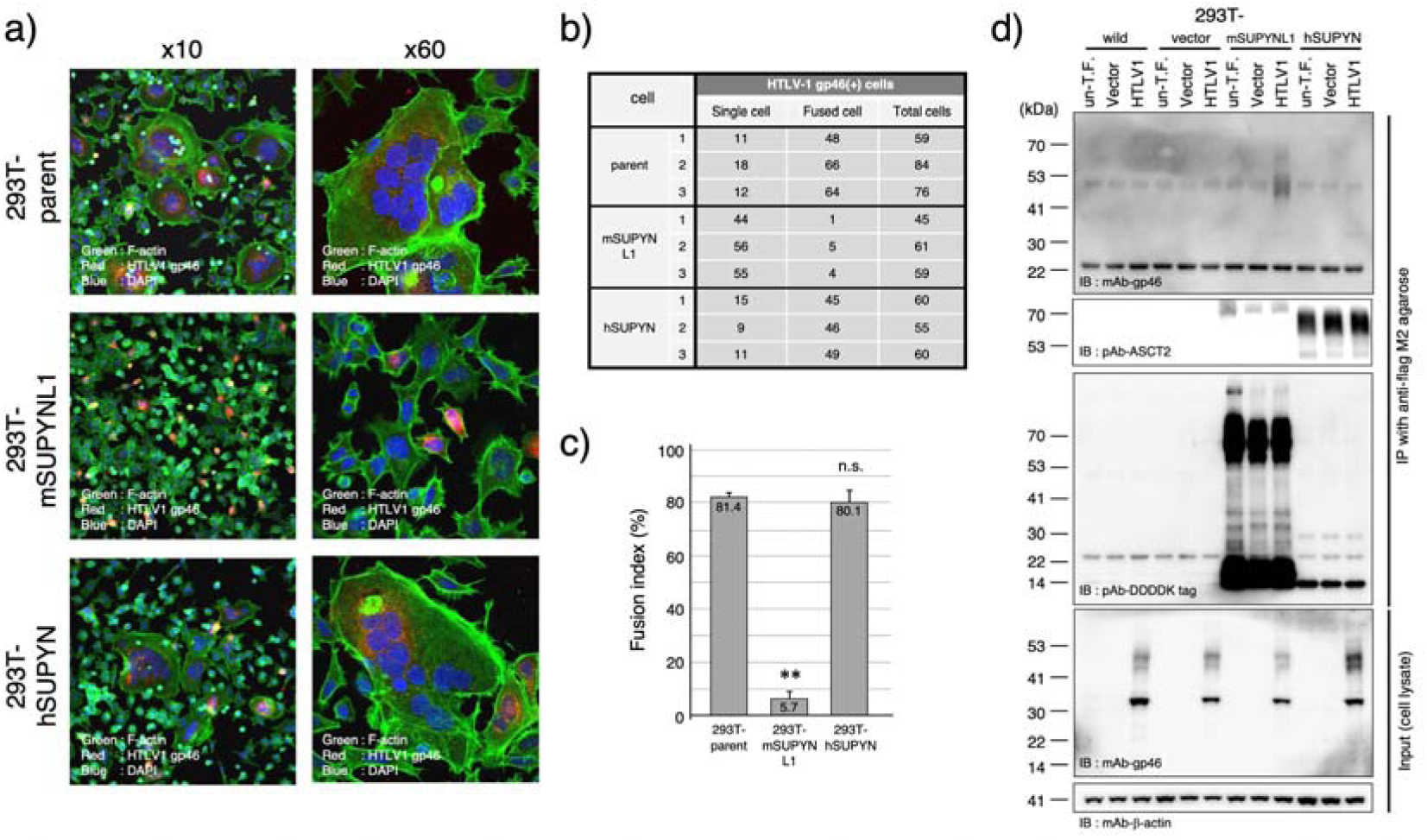
mSUPYNL1 suppresses HTLV-1 envelope-mediated membrane fusion through direct association with the SU glycoprotein gp46. a, Representative images of HTLV-1-induced syncytium formation. 293T cells stably expressing mSUPYNL1, human suppressyn (hSUPYN), or empty vector were transfected with the infectious HTLV-1 molecular clone pX1MT-M. Cells were stained for F-actin, HTLV-1 Env (gp46), and nuclei to visualize membrane fusion and viral Env expression. b, Quantification of multinucleated syncytia. Cells containing two or more nuclei were scored as syncytia. Three independent experiments were analyzed. c, Statistical analysis of HTLV-1-induced membrane fusion. Data represent mean ± s.d. from three independent experiments. Statistical significance was determined using two-tailed non-repeated measures ANOVA. **, p < 0.01; n.s., not significant. d, mSUPYNL1 directly associates with the HTLV-1 SU glycoprotein gp46. FLAG-tagged mSUPYNL1, hSUPYN, or empty vector was expressed in 293T cells together with HTLV-1 Env. Protein complexes were isolated by FLAG immunoprecipitation and analyzed by immunoblotting with anti-gp46 antibody. In parallel, interaction between hSUPYN and its known receptor ASCT2 was examined as a positive control. Input lysates demonstrate comparable protein expression, whereas immunoprecipitated fractions identify proteins directly associated with mSUPYNL1 or hSUPYN. Anti-FLAG and β-actin served as immunoprecipitation and loading controls, respectively. Uncropped immunoblots are shown in Supplementary Fig. 8.

To determine whether this inhibition reflected recognition of the viral envelope glycoprotein, we performed co-immunoprecipitation analyses using FLAG-tagged mSUPYNL1 and HTLV-1 Env. mSUPYNL1 specifically associated with the HTLV-1 surface glycoprotein gp46 (Fig. 3d), whereas hSUPYN showed no detectable association despite retaining its established interaction with ASCT2. These findings indicate that receptor-independent envelope glycoprotein recognition is unique to mSUPYNL1.

Together, these findings demonstrate that the mechanism underlying mSUPYNL1-mediated fusion suppression is not restricted to endogenous retroviral fusogens but extends to an exogenous pathogenic retrovirus. Thus, mSUPYNL1 recognizes envelope glycoproteins from both endogenous and exogenous retroviruses, identifying envelope glycoprotein recognition as a broad mechanism for suppressing retroviral membrane fusion independently of receptor competition.

### mSUPYNL1 is predominantly expressed in maternal decidual stromal and vascular-associated cells during placental development

To determine the physiological site of mSUPYNL1 expression during placental development, we generated a CRISPR/Cas9 knock-in/knockout mouse in which the coding region of mSUPYNL1 was replaced by an EGFP reporter cassette (Fig. 4a). This strategy enabled simultaneous disruption of mSUPYNL1 and visualization of endogenous transcriptional activity.

**Figure 4.**
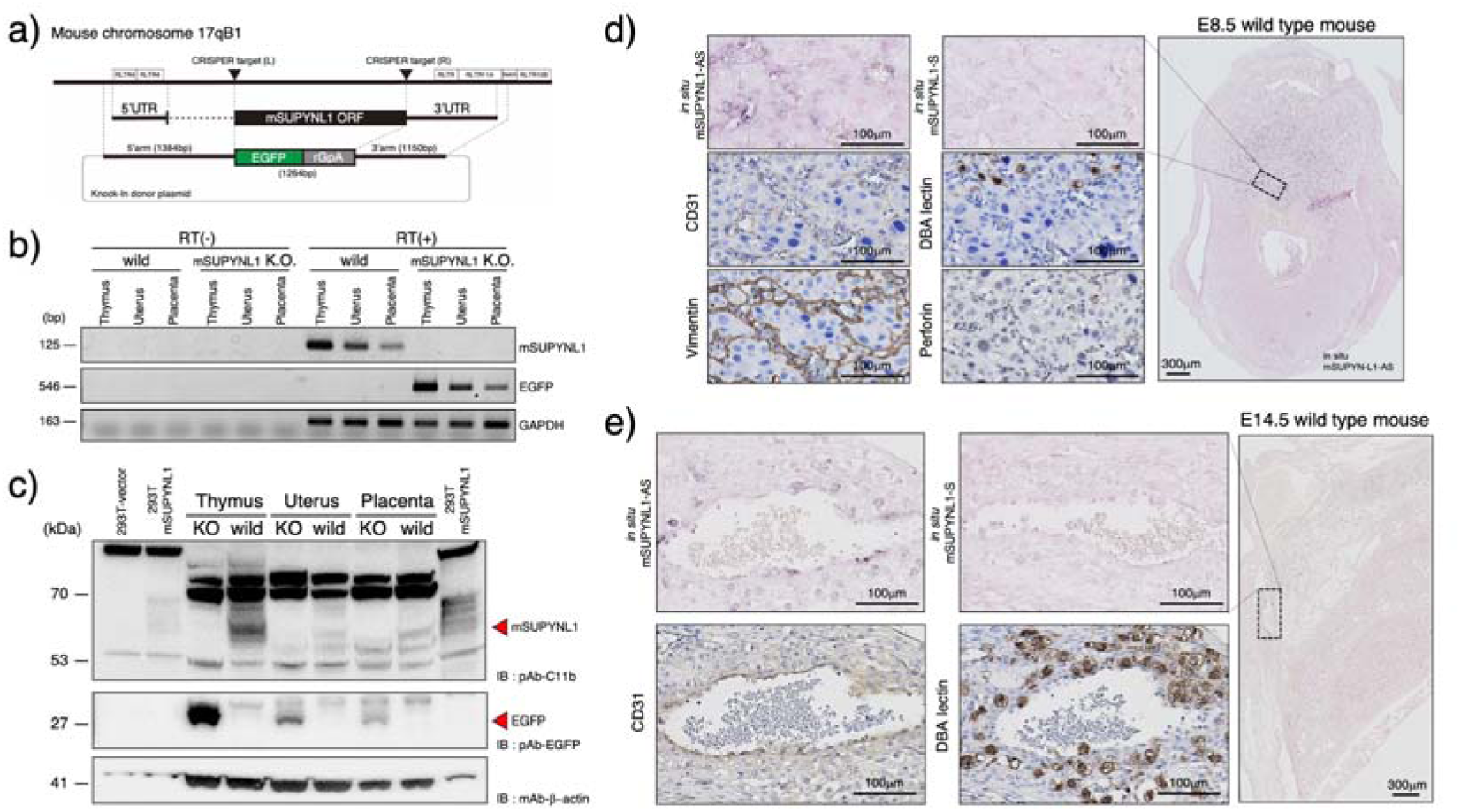
Generation of mSUPYNL1 knockout mice and characterization of mSUPYNL1 expression in vivo. a, Strategy for generation of the mSUPYNL1 knockout mouse. CRISPR/Cas9-mediated genome editing replaced the protein-coding region with an EGFP knock-in cassette. Positions of the two guide RNAs are indicated (L and R). rGpA, rabbit globin polyadenylation signal. b, RT-PCR analysis of mSUPYNL1 expression in wild-type and knockout mice. RNA isolated from thymus, uterus, and placenta was analyzed by RT-PCR. RT(-) controls lacking reverse transcriptase confirmed the absence of genomic DNA contamination. Primer sequences used for RT-PCR are listed in Supplementary Table 1. c, Detection of endogenous mSUPYNL1 protein by immunoblotting. Tissue lysates from wild-type and knockout mice were analyzed using anti-mSUPYNL1 (pAb-C11b) and anti-EGFP antibodies. Lysates from 293T cells expressing mSUPYNL1 or empty vector served as positive and negative controls, respectively. β-actin was used as a loading control. d, Localization of mSUPYNL1 transcripts in E8.5 implantation sites. In situ hybridization using sense and antisense probes was performed on serial sections, followed by immunohistochemistry for CD31, vimentin, DBA lectin, and perforin to identify mSUPYNL1-expressing cell populations associated with decidual vascular remodeling. e, Localization of mSUPYNL1 transcripts in E14.5 placentas. Serial sections were analyzed by in situ hybridization and immunostaining with CD31 and DBA lectin to characterize vascular endothelial and uNK-associated cell populations expressing mSUPYNL1. Uncropped gel images and immunoblots are provided in Supplementary Fig. 9.

RT-PCR and Northern blot analyses demonstrated broad mSUPYNL1 expression in multiple mouse tissues, including the placenta, spleen, thymus, kidney, and uterus (Fig. 4b and Supplementary Fig. 5a, b). EGFP expression faithfully recapitulated the endogenous transcription pattern, validating the reporter allele for subsequent localization studies.

Endogenous mSUPYNL1 protein was detected in the placenta, uterus, and spleen but was absent in knockout mice, whereas EGFP was readily detected from the targeted allele, confirming successful replacement of the endogenous locus (Fig. 4c).

In situ hybridization revealed that mSUPYNL1 expression was highly enriched in maternal decidual tissues during placental development. At embryonic day (E) 8.5, mSUPYNL1 transcripts were predominantly localized to decidual stromal cells surrounding maternal blood vessels (Fig. 4d and Supplementary Fig. 5c).

These signals closely overlapped vascular-associated stromal cell populations identified by CD31 and vimentin staining. Only limited overlap with DBA lectin-positive uterine natural killer (uNK) cells was observed, suggesting that decidual stromal cells constitute the principal site of mSUPYNL1 expression. EGFP in situ hybridization reproduced the endogenous expression pattern, confirming the fidelity of the reporter allele (Supplementary Fig. 5c). By E14.5, mSUPYNL1 expression became particularly prominent in vascular endothelial cells within both the decidua and uterine myometrium, as demonstrated by co-localization with CD31 (Fig. 4e and Supplementary Fig. 5d, e). Although mSUPYNL1-positive cells were frequently located adjacent to DBA lectin-positive uNK cells, direct co-localization remained limited. Comparable localization was observed using EGFP probes in knockout placentas. Expression was also detected in endothelial cells of the placental labyrinth, further supporting preferential expression in vascular-associated cell populations. Together, these findings identify the maternal decidua as the principal physiological site of mSUPYNL1 expression. The preferential localization of mSUPYNL1 to decidual stromal and vascular-associated cells contrasts sharply with the trophoblast-enriched expression of

hSUPYN and suggests that ERV-derived fusion suppressors have undergone functional diversification during mammalian evolution.

## Discussion

In this study, we show that mouse suppressyn-like 1 (mSUPYNL1) functions as an endogenous retrovirus (ERV)-derived envelope protein that inhibits membrane fusion through a mechanism fundamentally distinct from that used by hSUPYN. Whereas hSUPYN suppresses syncytin-1 via receptor interference^20,22–24^, mSUPYNL1 instead binds the surface (SU) glycoproteins of multiple retroviral envelope proteins, thereby inhibiting membrane fusion independently of receptor usage. Together, these observations identify direct envelope glycoprotein recognition as an evolutionarily distinct mechanism of ERV-derived fusion suppression and suggest that receptor-independent envelope recognition provides an alternative molecular strategy for controlling trophoblast fusion.

One of the most striking features of mSUPYNL1 is its broad reactivity toward retroviral envelope proteins. In addition to inhibiting both murine and human syncytins, mSUPYNL1 also interacts with the SU glycoprotein (gp46) of the exogenous retrovirus, HTLV-1, and suppresses Env-dependent syncytium formation^25,26^. Combined with the co-immunoprecipitation data, these results support the idea that mSUPYNL1 recognizes conserved structural determinants within retroviral envelope glycoproteins rather than specific cellular retroviral receptors. Such a mechanism could account for its ability to inhibit membrane fusion driven by phylogenetically diverse envelope proteins despite their use of distinct entry receptors. Whether mSUPYNL1 recognizes a common structural motif or multiple independent epitopes shared among these envelope proteins remains an important question for future structural studies.

The ability of mSUPYNL1 to inhibit HTLV-1 Env-mediated syncytium formation extends the potential significance of this mechanism beyond endogenous placental fusogens. Unlike hSUPYN, mSUPYNL1 binds the viral envelope glycoprotein gp46, raising the possibility that recognition of newly synthesized envelope proteins within the cell could restrict subsequent cell-to-cell viral transmission. Although the physiological importance of this antiviral activity is still unclear, our results suggest that domesticated ERV-derived envelope proteins may have been co-opted not only to regulate physiological membrane fusion but also to limit membrane fusion mediated by exogenous retroviruses.

The tissue distribution of mSUPYNL1 further differentiates it from hSUPYN and supplies insights into its physiological role. Whereas hSUPYN expression is largely restricted to trophoblast populations in the human placenta^22,27^, mSUPYNL1 is broadly expressed, with prominent expression in decidual stromal cells, vascular endothelial cells, and hematopoietic tissues. Its enrichment at the maternal–fetal interface raises the possibility that receptor-independent regulation of membrane fusion contributes to coordinated interactions among trophoblasts, stromal cells, endothelial cells, and immune cells during placental morphogenesis and vascular remodeling^28,29^. Whether mSUPYNL1 directly controls these cellular interactions remains to be determined, and studies using mSUPYNL1-deficient mice should provide an opportunity to address this question in vivo.

The evolutionary origins of hSUPYN and mSUPYNL1 provide an interesting example of functional convergence. Human SUPYN and mSUPYNL1 most likely originated from independent ERV integration events during primate and rodent evolution, respectively, yet both acquired the capacity to suppress membrane fusion through different molecular mechanisms. This pattern implies that evolutionary selection acted on the biological function -fusion suppression- while permitting distinct biochemical solutions to emerge independently. As opposed to representing a single evolutionary strategy, these proteins illustrate that ERV-derived envelope genes can be adapted through multiple molecular mechanisms to regulate membrane fusion.

Our study also has several limitations. Although our expression analyses strongly implicate mSUPYNL1 in placental biology, its physiological role in vivo is still incompletely understood. In particular, further analyses of mSUPYNL1-deficient mice will be important for determining how loss of this protein affects placental development and fetal growth. Likewise, structural studies of mSUPYNL1 in complex with retroviral envelope glycoproteins will be needed to define the molecular basis and full extent of its broad envelope recognition and receptor-independent inhibition of membrane fusion. Taken together, our data identify mSUPYNL1 as the first ERV-derived envelope protein shown to suppress membrane fusion through direct recognition of retroviral envelope glycoproteins rather than receptor interference. By uncovering this mechanism,

our study expands the range of molecular strategies through which domesticated retroviral envelope proteins regulate membrane fusion. The mSUPYNL1 mouse model serves as a valuable experimental platform to investigate how ERV-derived fusion suppressors contribute to placental biology, antiviral defense, and the evolution of ERV-derived host functions.

## Methods

### Identification of candidate endogenous retroviral envelope genes from mouse placental RNA-seq datasets

To identify endogenous retrovirus (ERV)-derived envelope genes expressed in the mouse placenta, publicly available RNA-seq data generated from mouse placental tissue were analyzed. Mouse Placenta RNA-seq datasets were downloaded from the NCBI Sequence Read Archive (SRA accession numbers SRR392616 and SRR392617). Adapter sequences and low-quality bases were trimmed using Cutadapt v1.8.3^30^. The processed reads were aligned to the mouse reference genome (GRCm38.p1) using TopHat2 v2.1.1^31^. Transcript abundance was then quantified with Cufflinks v2.2.1^32^ using the mouse protein-coding endogenous viral element (EVE) annotation data (Mmus38.geve.m_v1.gtf) obtained from the gEVE database^21^.

Candidate ERV loci were ranked according to transcript abundance in placental tissue. Genomic sequences corresponding to highly expressed candidates were subsequently examined for intact open reading frames (ORFs) capable of encoding putative envelope proteins. Based on transcript abundance and ORF integrity, fifteen candidate ERV-derived *env* loci were selected for subsequent molecular cloning and functional characterization.

### Cell culture

Human embryonic kidney 293T cells (ATCC CRL-3216) and Madin–Darby canine kidney (MDCK) cells (ATCC CCL-34) were obtained from the American Type Culture Collection (ATCC, Manassas, VA, USA). 293T cells were maintained in Dulbecco’s Modified Eagle Medium, whereas MDCK cells were cultured in Eagle’s Minimum Essential Medium. Both media were supplemented with 10% heat-inactivated fetal bovine serum (FBS; Thermo Fisher Scientific, Waltham, MA, USA). Cells were maintained at 37°C in a humidified atmosphere containing 5% CO[.

### Plasmid construction

Full-length mSUPYNL1, candidate ERV-derived genes (C7, C14, and C15), syncytin-A, and syncytin-B cDNAs were amplified by RT-PCR from mouse placental RNA. PCR products were subsequently re-amplified using primers containing appropriate restriction enzyme recognition sites and cloned into the pGEM-T Easy vector (A1360: Promega, WI, USA) for sequence verification. Primer sequences used for cloning are listed in Supplementary Table 1. The nucleotide sequences of candidate genes C7, C14, and C15 have been omitted because these genes are currently under independent investigation and will be described in a separate study. Only clones exhibiting 100% sequence identity with the corresponding reference sequences were used for subsequent experiments. The coding sequences verified by Sanger sequencing, of mSUPYNL1 (GenBank accession AK088743), syncytin-A (NM_001013751), and syncytin-B (NM_173420.3) were subcloned into the mammalian expression vector pFlag-EF1either in-frame with a C-terminal FLAG epitope tag for protein expression and immunoblotting or without an epitope tag for cell fusion assays^20^. Expression constructs for human syncytin-1 and syncytin-2 have been described previously and were used in this study without modification. To investigate protein–protein interactions, DNA fragments encoding the surface (SU) and transmembrane (TM) subunits of mSUPYNL1, syncytin-A, syncytin-B, syncytin-1, and syncytin-2 were amplified using restriction site-containing primers and cloned into pFlag-EF1-derived expression vectors carrying either a C-terminal FLAG tag or a hemagglutinin (HA) tag, as indicated in each experiment. For the generation of stable mammalian cell lines, the full-length mSUPYNL1 coding sequence was additionally subcloned into the pCAG mammalian expression vector under the control of the CAG promoter^20^.

### Antibodies

A rabbit polyclonal antibody against mSUPYNL1 (designated pAb-C11b) was generated by immunizing rabbits with a synthetic peptide corresponding to the mSUPYNL1 amino acid sequence (Cys-QELRPRSGPWDLLKA-OH). The antibody was affinity-purified using a peptide-conjugated affinity column (Scrum Inc., Tokyo, Japan) and used for immunoprecipitation and immunoblotting experiments.

A complete list of commercially available primary and secondary antibodies used for immunoprecipitation, western blotting, immunohistochemistry, and immunofluorescence staining is provided in Supplementary Table 2.

### Immunoprecipitation and western blotting

Mouse tissues (thymus, uterus, and placenta) and cultured cells were lysed in RIPA buffer containing 50 mM Tris-HCl (pH 8.0), 150 mM NaCl, 0.5% sodium deoxycholate, 0.1% SDS, and 1% NP-40 substitute supplemented with a protease inhibitor cocktail (161-26023: FUJIFILM Wako Pure Chemical Corporation, Osaka, Japan). Protein lysates were clarified by centrifugation and quantified before analysis. Protein concentrations were determined using BCA assays (T9300A: Takara, Shiga, Japan).

For FLAG-mediated immunoprecipitation, 300 μg of total protein was incubated with anti-FLAG M2 agarose (A2220: Sigma-Aldrich, Darmstadt, Germany) according to the manufacturer’s instructions. Following extensive washing, immunoprecipitated proteins were subjected to SDS-PAGE and immunoblot analysis.

To detect secreted mSUPYNL1 protein, conditioned culture medium was supplemented with protease inhibitor cocktail immediately after collection. The medium was incubated with 0.5 μg of affinity-purified pAb-C11b for 2 h at 4°C, followed by the addition of 10 μL Protein G agarose beads (sc-2002: Santa Cruz Biotechnology, Texas, USA) and further incubation overnight at 4°C with gentle rotation. Immunocomplexes were recovered by centrifugation, washed extensively, and analyzed by western blotting. Protein samples were separated by conventional SDS-PAGE or by NuPAGE 4–12% Bis-Tris precast gels (NP0322BOX and NW04120BOX: Thermo Fisher Scientific, MA, USA ) according to the manufacturer’s protocol. Proteins were transferred onto PVDF membranes using the iBlot 2 Dry Blotting System (IB21001: Thermo Fisher Scientific, MA, USA). Membranes were incubated with the appropriate primary antibodies followed by horseradish peroxidase (HRP)-conjugated goat anti-rabbit IgG (7074: Cell Signaling Technology, MA, USA) or HRP-conjugated goat anti-mouse IgG (7076: Cell Signaling Technology, MA, USA). Immunoreactive signals were visualized using Chemi-Lumi One Super (0230-14: Nacalai Tesque, Kyoto, Japan) according to the manufacturer’s instructions.

### Generation of mSUPYNL1 knockout mice

The mSUPYNL1 knockout mouse line was generated by CRISPR/Cas9-mediated genome editing on a C57BL/6J genetic background. Two guide RNAs targeting the 5′ and 3′ regions flanking the mSUPYNL1 coding sequence were designed and cloned individually into the pX330 CRISPR/Cas9 expression vector (#42230: Addgene, MA, USA). The guide RNA sequences were as follows: left guide, 5′-TGAAGTCTCGATCACTAGAC-3′; right guide, 5′-CGTATTAAAATGCTAACCCT-3′.

A donor plasmid (pDonor-D17-EGFP) carrying the EGFP reporter cassette flanked by homologous genomic sequences was used to replace the endogenous mSUPYNL1 locus by homology-directed repair. Plasmid DNA was purified using the FastGene Plasmid Mini Kit (FG-90402: Nippon Genetics, Tokyo, Japan) and mixed at final concentrations of 5 ng/μl for each CRISPR/Cas9 vector and 10 ng/μl for the donor plasmid. Primer sequences used for cloning are listed in Supplementary Table 1.

The plasmid mixture was filtered through a 0.22-μm membrane filter and microinjected into the pronuclei of fertilized C57BL/6J zygotes. Surviving embryos were transferred into the oviducts of pseudo pregnant ICR recipient females using standard embryo transfer procedures. Founder mice were screened by genomic PCR using DNA isolated from tail biopsies, and targeted alleles were confirmed by sequencing. Genotyping of subsequent generations was performed by PCR using genomic DNA extracted from tail samples. Successful replacement of the endogenous mSUPYNL1 coding region with the EGFP cassette is illustrated in Fig. 3a. Homozygous knockout mice were viable and were used for all subsequent expression and phenotypic analyses.

### HTLV-1-induced cell fusion assay

Control 293T cells, 293T cells stably expressing mSUPYNL1, and 293T cells stably expressing hSUPYN were seeded into 12-well plates at 1.8 × 10[ cells per well. Twenty-four hours later, cells were transiently transfected with 1 μg of the infectious HTLV-1 molecular clone pX1MT-M^33^ using Lipofectamine 3000 (L3000015: Thermo Fisher Scientific, MA, USA) according to the manufacturer’s instructions. At 24 h post-transfection, cells were detached with 0.25% trypsin-EDTA, replated onto BioCoat Poly-D-Lysine four-well culture slides (354577: Corning, NY, USA) at 1 × 10[ cells per well, and cultured for an additional 24 h. Cells were then fixed with 2% paraformaldehyde, permeabilized with 0.2% Triton X-100, and incubated with the rat monoclonal anti-HTLV-1 Env antibody LAT-27^34^, followed by Alexa Fluor 568-conjugated goat anti-rat IgG (A-11077: Thermo Fisher Scientific, MA, USA). F-actin and nuclei were visualized using Alexa Fluor 488-conjugated phalloidin (A-12379: Thermo Fisher Scientific, MA, USA) and DAPI, respectively. Fluorescence images were acquired using an FV3000 confocal laser-scanning microscope (Olympus, Tokyo, Japan) and processed using FV31S-SW software (Olympus, Tokyo, Japan). Cells containing two or more nuclei within a continuous plasma membrane were scored as syncytia. At least 40 cells were analyzed per experimental condition.

### Co-immunoprecipitation of mSUPYNL1 with HTLV-1 envelope protein

Parental 293T cells, vector control cells, 293T cells stably expressing mSUPYNL1, and 293T cells stably expressing hSUPYN were seeded into 12-well plates at 2 × 10[ cells per well. Cells were transiently transfected with 2 μg of either the empty control vector (pCAG-GS-vec) or the HTLV-1 envelope expression plasmid (pCAG-GS-HTLV1env)^35^ using Lipofectamine 2000 (11668027: Thermo Fisher Scientific, MA, USA). The HTLV-1 envelope expression plasmid was kindly provided by Dr. Kenta Tezuka (National Institute of Infectious Diseases, Japan Institute for Health Security). Forty-eight hours after transfection, cells were harvested and lysed in RIPA buffer as described above. FLAG-tagged proteins were immunoprecipitated using anti-FLAG M2 agarose. Immunoprecipitated proteins were separated by SDS-PAGE and analyzed by western blotting using an anti-HTLV-1 gp46 antibody (sc-53890: SantaCruz, TX, USA) followed by TrueBlot ULTRA anti-mouse Ig HRP (18-8817-30: ROCKLAND, PA, USA) to detect the interaction between mSUPYNL1 and the HTLV-1 envelope glycoprotein.

### In situ hybridization

Digoxigenin (DIG)-labeled antisense and sense RNA probes for mSUPYNL1 and EGFP were synthesized from pGEM-T Easy plasmids containing the corresponding cDNA fragments (mSUPYNL1, 501bp fragment; EGFP, 720bp fragment) using the DIG RNA Labeling Kit (SP6/T7) (11175025910: Roche Diagnostics, Mannheim, Germany) according to the manufacturer’s instructions. Paraffin-embedded uteroplacental tissues were sectioned at 6 μm. Tissue sections were deparaffinized, rehydrated, post-fixed in 10% neutral buffered formalin for 30 min at 37°C, treated with 0.2% HCl for 10 min, and digested with 5 μg/mL proteinase K for 10 min at 37°C. Subsequent hybridization and washing procedures were performed using the Genostaff in situ hybridization kit (SRK-02: Genostaff, Tokyo, Japan) according to the manufacturer’s protocol. Hybridization was carried out overnight at 60°C in G-Hybo-L hybridization buffer containing 1 μg/mL DIG-labeled RNA probe. Following high-stringency washes with 1× and 0.1× G-Wash buffer at 60°C, hybridized probes were detected using alkaline phosphatase-conjugated anti-digoxigenin antibody (148-06211, FUJIFILM Wako, Osaka, Japan) diluted 1:2,000. Color development was performed using NBT/BCIP substrate at 4°C overnight. Sections were counterstained with Kernechtrot, mounted using G-Mount, and Images were acquired using a BZ-X710 digital microscope (Keyence, Osaka, Japan) and processed using BZ-X Analyzer software.

## Acknowledgements

We thank Dr. Seiya Mizuno (Institute of Medicine, University of Tsukuba) for generating the mSUPYNL1 knockout mouse line using CRISPR/Cas9 genome-editing technology. We are grateful to Dr. Kenta Tezuka (National Institute of Infectious Diseases, Japan Institute for Health Security) for providing the pCAG-GS-HTLV1env expression plasmid used in the HTLV-1 Env interaction studies. We also thank Prof. Masao Matsuoka and Prof. Jun-ichiro Yasunaga (Kumamoto University) for providing the HTLV-1 molecular clone pX1MT-M, and Prof. Yuetsu Tanaka (Faculty of Medicine, University of the Ryukyus) for providing the HTLV-1-specific antibody used in this study. We are grateful to Mako Morioka and Kana Kurosaki (School of Medicine, Hiroshima University) for their technical assistance in constructing the syncytin expression plasmids. We also thank Yoko Hayashi and the staff of the Center for Life Science, Natural Science Center for Basic Research and Development, Hiroshima University, for their expert assistance with flow cytometric analyses and DNA sequencing. The affiliations listed above correspond to those at the time the work was performed and may have changed since then. This manuscript was edited for English language and conciseness with the assistance of ChatGPT (OpenAI). A preprint version of this manuscript has been deposited in bioRxiv (bioRxiv 2026.07.22.739964; doi: https://doi.org/10.64898/2026.07.22.739964).

## Author contributions

J.S. and D.J.S. contributed equally to this work.

J.S., D.J.S., and Y.K. conceived and designed the study.

J.S. and M(a).S. performed molecular and cellular experiments and analyzed the data.

S.N. performed the bioinformatic analyses.

M.H. and M.S. carried out HTLV-1 infection experiments and associated analyses. Y.S., H.T., and J.S. generated and characterized the mouse models and performed the in vivo analyses.

J.S., D.J.S., T.N., Y.J., and Y.K. interpreted the data.

J.S. and D.J.S. wrote the manuscript with input from all authors. All authors reviewed and approved the final manuscript.

## Additional information

### Data availability

The data that support the findings of this study are available from the corresponding author upon reasonable request.

### Data and Code Availability

The nucleotide and protein sequence of mouse suppressyn-like 1 (mSUPYNL1) has been deposited in GenBank under accession number PZ682522.

