## Supplementary material for "Mouse suppressyn-like 1 suppresses membrane fusion through envelope glycoprotein recognition": supplymentary figures

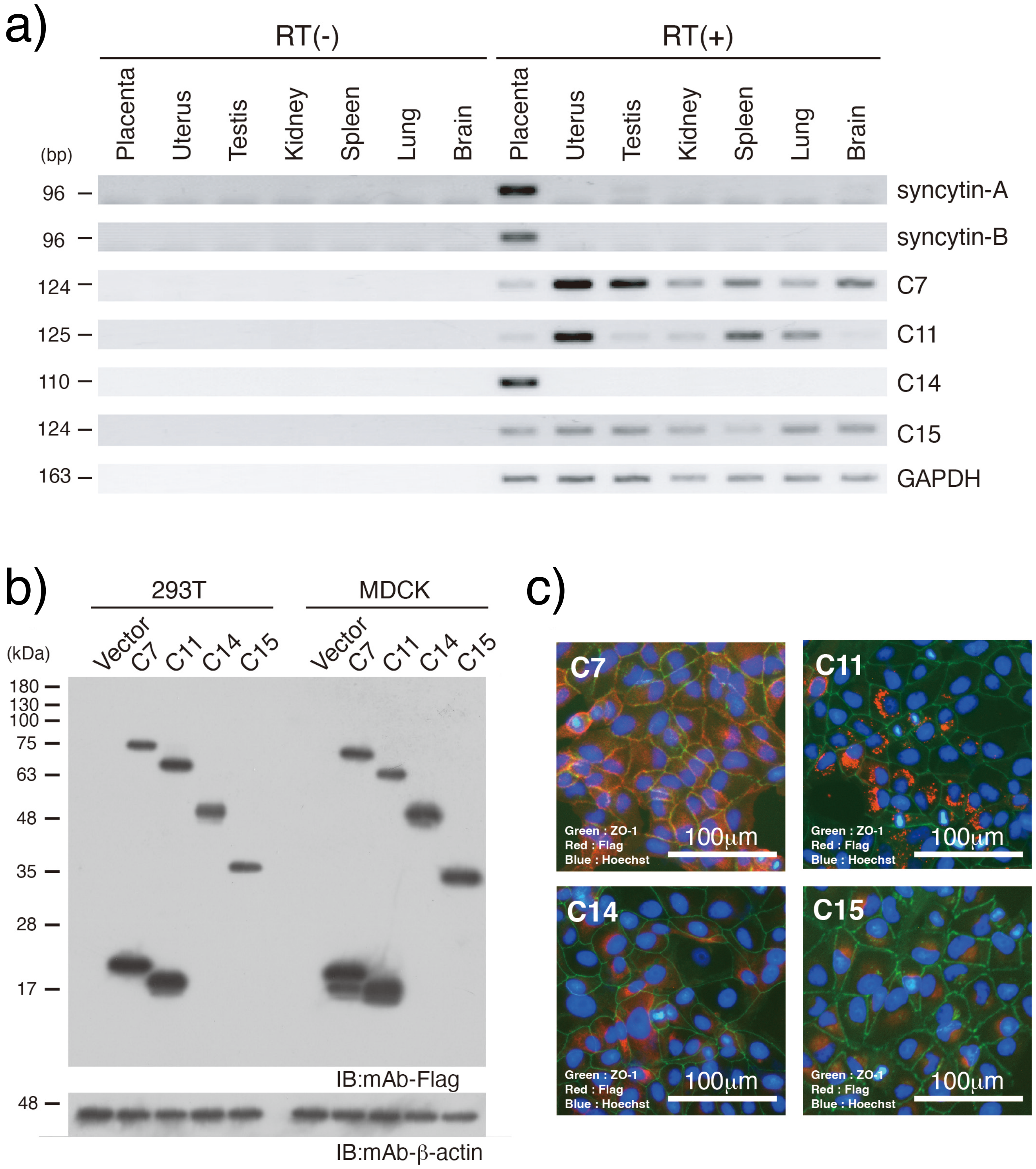


**Supplementary Figure 1. Isolation and characterization of candidate mouse suppressyn proteins.**

a. Expression analysis of candidate genes in various mouse tissues by RT-PCR. Syncytin-A and syncytin-B were used as placental expression controls, and mouse β-actin was used as an internal control. b. The predicted open reading frames (ORFs) of four candidate genes were cloned and stably expressed in 293T and MDCK cells, and the corresponding proteins were detected by immunoblotting. c. Subcellular localization of the translated products of the four candidate genes in MDCK cells was examined by immunocytochemistry. Candidate proteins were expressed as C-terminally Flag-tagged fusion proteins and are shown in red. Cell boundaries were visualized by ZO-1 immunostaining (green), and nuclei were stained with Hoechst dye (blue).

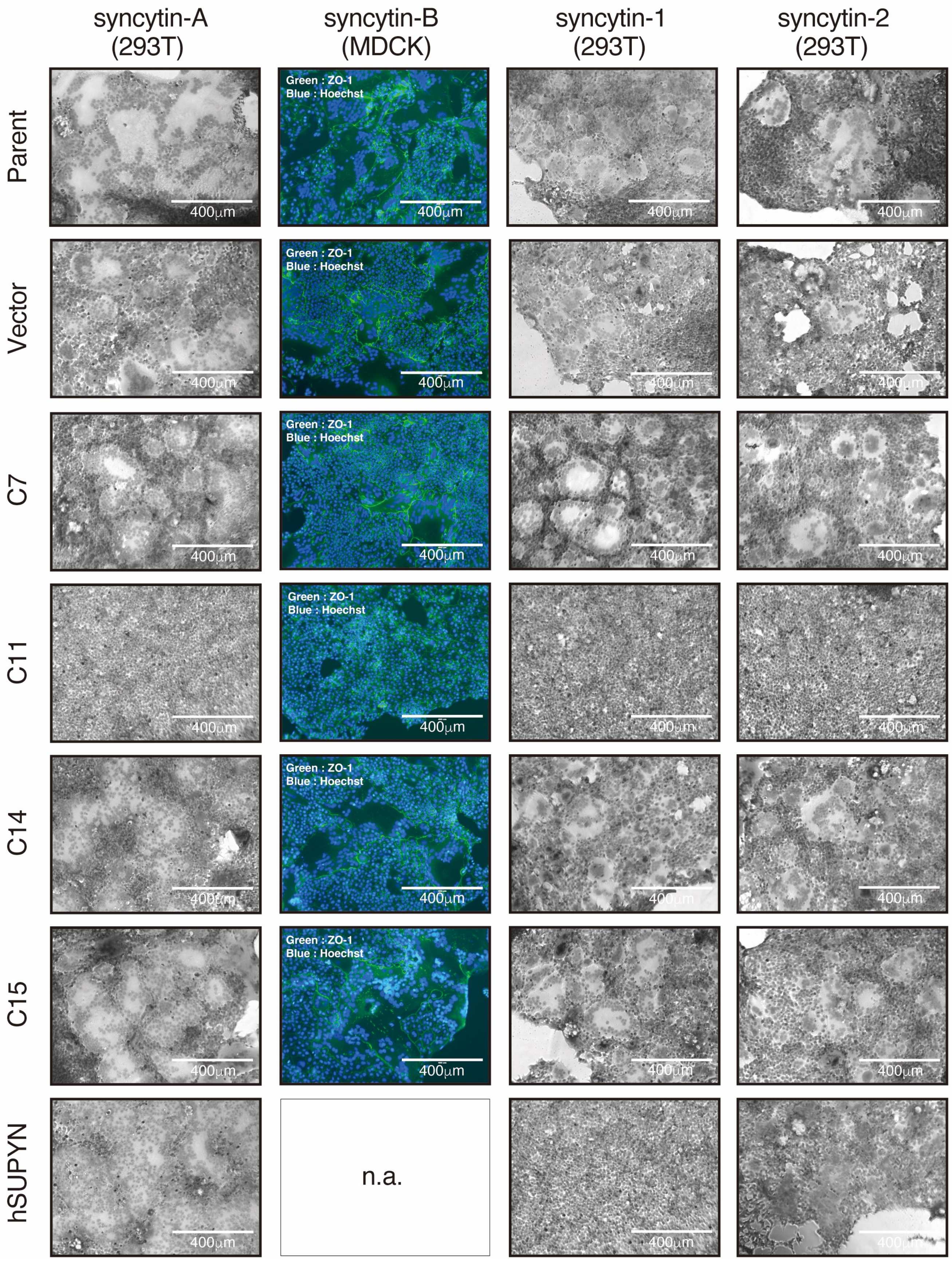


**Supplementary Figure 2. Evaluation of the inhibitory effects of four candidate proteins on syncytin-mediated cell fusion.**

The four candidate genes (C7, C11, C14, and C15), together with human suppressyn (hSUPYN), were stably expressed in 293T cells (for syncytin-A, syncytin-1, and syncytin-2 assays) or MDCK cells (for syncytin-B assays). Syncytin-mediated cell fusion was evaluated by transient expression of each syncytin in these cells, followed by hematoxylin staining or immunostaining of cell boundaries with an anti-ZO-1 antibody. The inhibitory activity of hSUPYN was evaluated only in the 293T cell-based syncytin fusion assay, as 293T cells are susceptible to syncytin-1-mediated fusion.

```
mSUPYNL1  51   SPMPHEPHNLTWVITNTATGTVINSTSHLAPIGTWFPDLVFDLCALADGTWDPLGLV---  107
               +P PH+ +N+TW ITN  TGT  N+TS L  +   FP L FDLC +   TW+P G
FeLV env  2    NPSPHQVYNITWTITNLVTGTKANATSMLGTLTDAFPTLYFDLCDIIGNTWNPSGQEPFP  61

mSUPYNL1  108  -WGCQHPAQHSELRRTPFYVCPGEGRSPQQTATCGGPESYFCAYWGCESTGSISWTPPIK  166
                +GC  P +    R T FYVCPG     Q    CGGP+  FCA WGCE+TG   W P
FeLV env  62   GYGCDQPMRRWRQRNTAFYVCPGHANQKQ----CGGPQDGFCAVWGCETTGEAYWKPTSS  117

mSUPYNL1  167  DDLIKVQRSPGS-----ESQGW-GYGLQPKLDANDLGGPCSSNCNPITVQFTPKGKESVG  220
                D I V++            GW G      + ++  G      CNP+ +QFT KG+++
FeLV env  118  WDYITVKKGVTQGIYQCSGGGWCGPCYDKAVHSSTTGASEGGRCNPLILQFTQKGRQT-S  176

mSUPYNL1  221  WEKGKTWGLRLYVSGYDYGVMFTIQLSARQGMP---LGPNPALPPVGLTVTQSQITVPIS  277
               W+  K+WGLRLY SGYD   +F++        P   +GPN  LP       QSQI   ++
FeLV env  177  WDGPKSWGLRLYRSGYDPIALFSVSRQVMAITPPQAMGPNLVLPDQKPPSRQSQIESKVA  236

mSUPYNL1  278  QELRPR------SGPW-----------------DLLKATYLVLNNSKPELTSKCWLCLDA  314
                +  PR      SGP                   L++ TYL LN + P  T  CWLCL +
FeLV env  237  TQ-SPRRNTSSVSGPGTPTTISPKRTGTGDRLISLIQGTYLALNATNPNKTKDCWLCLVS  295

mSUPYNL1  315  QPPYYEGIAVPGNYTTSTDNK-DCRWQYPGKGRLTLELIQGKGLCFGTIPHTHRHLCNSI  373
               +PPYYEGIA+ GNY+  T+    C      + +LT+  + G+GLC GT+P TH+ LCN
FeLV env  296  RPPYYEGIAILGNYSNQTNPPPSCLSTL--QHKLTISEVSGQGLCIGTVPKTHQALCNKT  353

mSUPYNL1  374  QEAPDRDAYIIPPINTWWACTTGLTSCIHTQALNLTNGFCVLVQLAPRILRYSDE-MGTR  432
               Q+      Y+  P  T+WAC TGLT CI    LN T+ FCVL++L PR+  +  E + T
FeLV env  354  QQGHTGAHYLAAPNGTYWACNTGLTPCISMAVLNWTSDFCVLIELWPRVTYHQPEYVYTH  413

mSUPYNL1  433  LSPFPAARAKRA----TVAEILGASAFD-------------FQDENYKVLSSAIDADLME  475
                     AR++R     TVA +LG                    +   ++ L  A+  D+
FeLV env  414  FD--KTARSRREPISLTVALMLGGLTVGGIAAGVGTGTKALLETAQFRQLQMAMHTDIQA  471

mSUPYNL1  476  LGKSLSKSKTSLTSLREATLRNQREQDFQSLQQDGLCKPLEKRCCTFVDNLKHARELLAK  535
               L +S+S  + SLTSL E  L+N+R  D   LQ  GLC  L++ CC + D+    R+ +AK
FeLV env  472  LEESVSALEKSLTSLSEVVLQNRRGLDILFLQGGGLCAALKEECCFYADHTGLVRDSMAK  531

mSUPYNL1  536  VRKRLEDRKREMEQGQ----------PVWITVIATLSG  563
               +R+RL+ R++  +  Q          P + T+I+++ G
FeLV env  532  LRERLKQRQQLFDSQQGWFEGWFNRSPWFTTLISSIMG  569
```

**Supplementary Figure 3. BLASTp analysis of mSUPYNL1.**

BLASTp analysis revealed sequence similarity between mSUPYNL1 and the envelope protein of feline leukemia virus (FeLV; partial sequence, BAK41678.1, length 612 amino acids). mSUPYNL1 exhibited 36% amino acid sequence identity with the FeLV envelope protein.

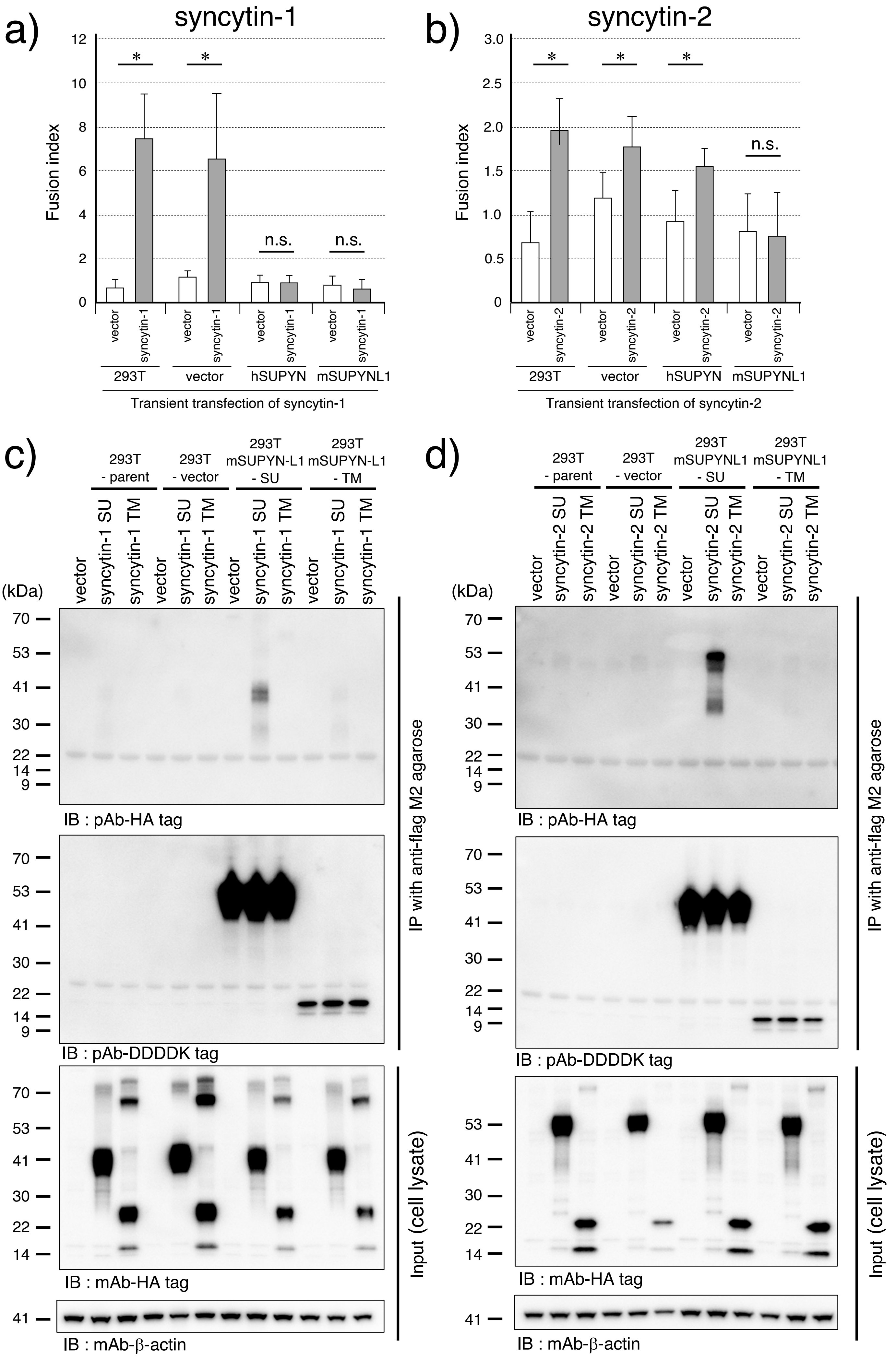


**Supplementary Figure 4. Inhibitory effects of mSUPYNL1 on syncytin-1- and syncytin-2-mediated cell fusion and identification of interacting proteins.**

a, b. Similar to Figure 2b, 293T cells stably expressing mSUPYNL1 were transiently transfected with syncytin-1 (100 ng) or syncytin-2 (500 ng) expression plasmids, and syncytin-mediated cell fusion was quantified by flow cytometry. Cells expressing hSUPYN were analyzed in parallel as a control. The fusion activity was statistically compared with that of cells transiently expressing the corresponding empty vector using the Mann–Whitney U test. **, $p < 0.01$; n.s., not significant.

c, d. To identify interacting proteins, the SU and TM subunits of mSUPYNL1 were individually expressed in 293T cells, together with the SU or TM subunits of syncytins. Co-immunoprecipitation assays were performed using an anti-Flag antibody to detect Flag-tagged mSUPYNL1-derived proteins and associated syncytin proteins.

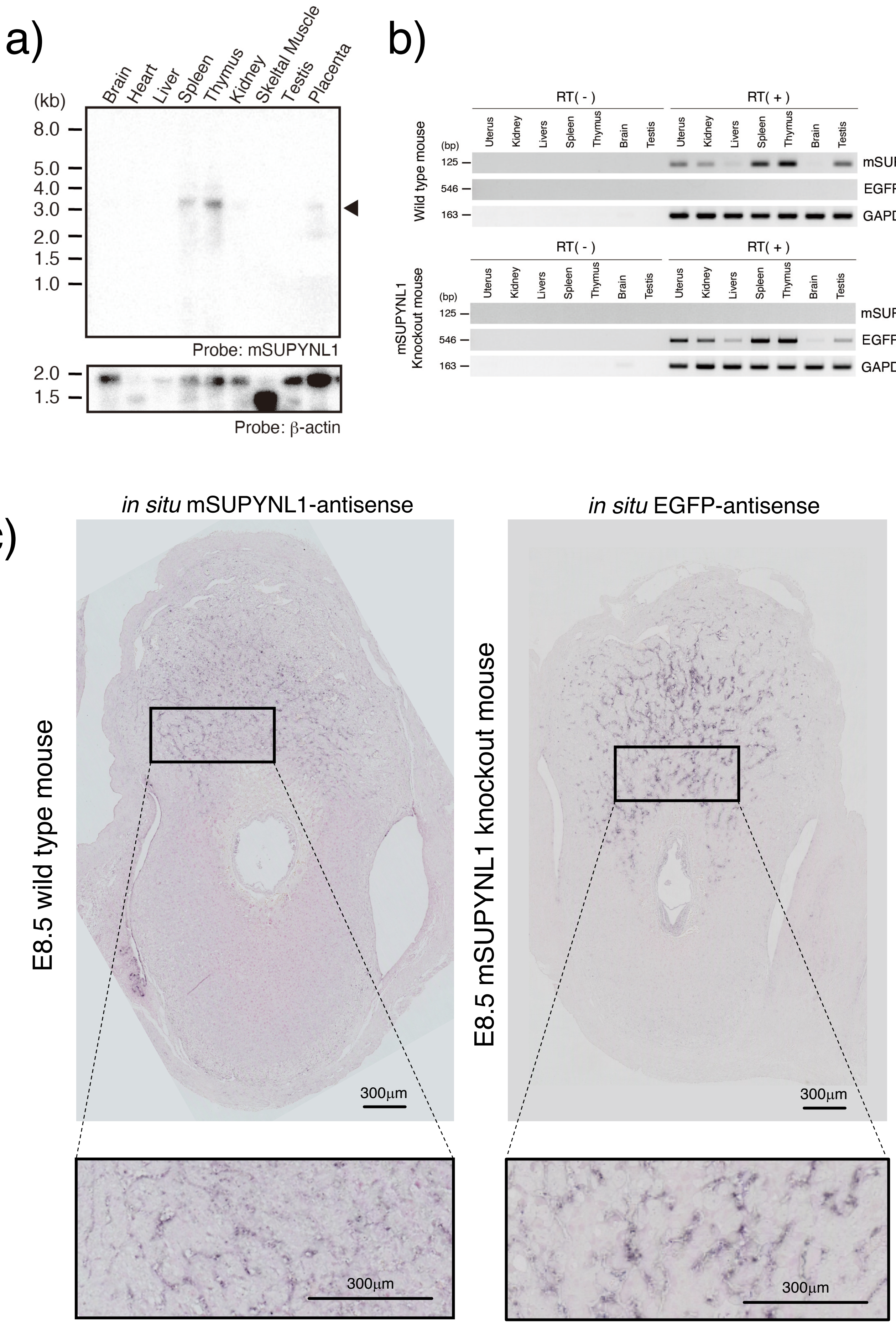

a)
Brain
Heart
Liver
Spleen
Thymus
Kidney
Skeltal Muscle
Testis
Placenta
(kb)
8.0
5.0
4.0
3.0
2.0
1.5
1.0
Probe: mSUPYNL1
2.0
1.5
Probe: β-actin
b)
RT( - )
RT( + )
Wild type mouse
(bp)
Uterus
Kidney
Livers
Spleen
Thymus
Brain
Testis
125
546
163
mSUPYNL1
EGFP
GAPDH
mSUPYNL1 Knockout mouse
c)
in situ mSUPYNL1-antisense
in situ EGFP-antisense
E8.5 wild type mouse
E8.5 mSUPYNL1 knockout mouse
300µm

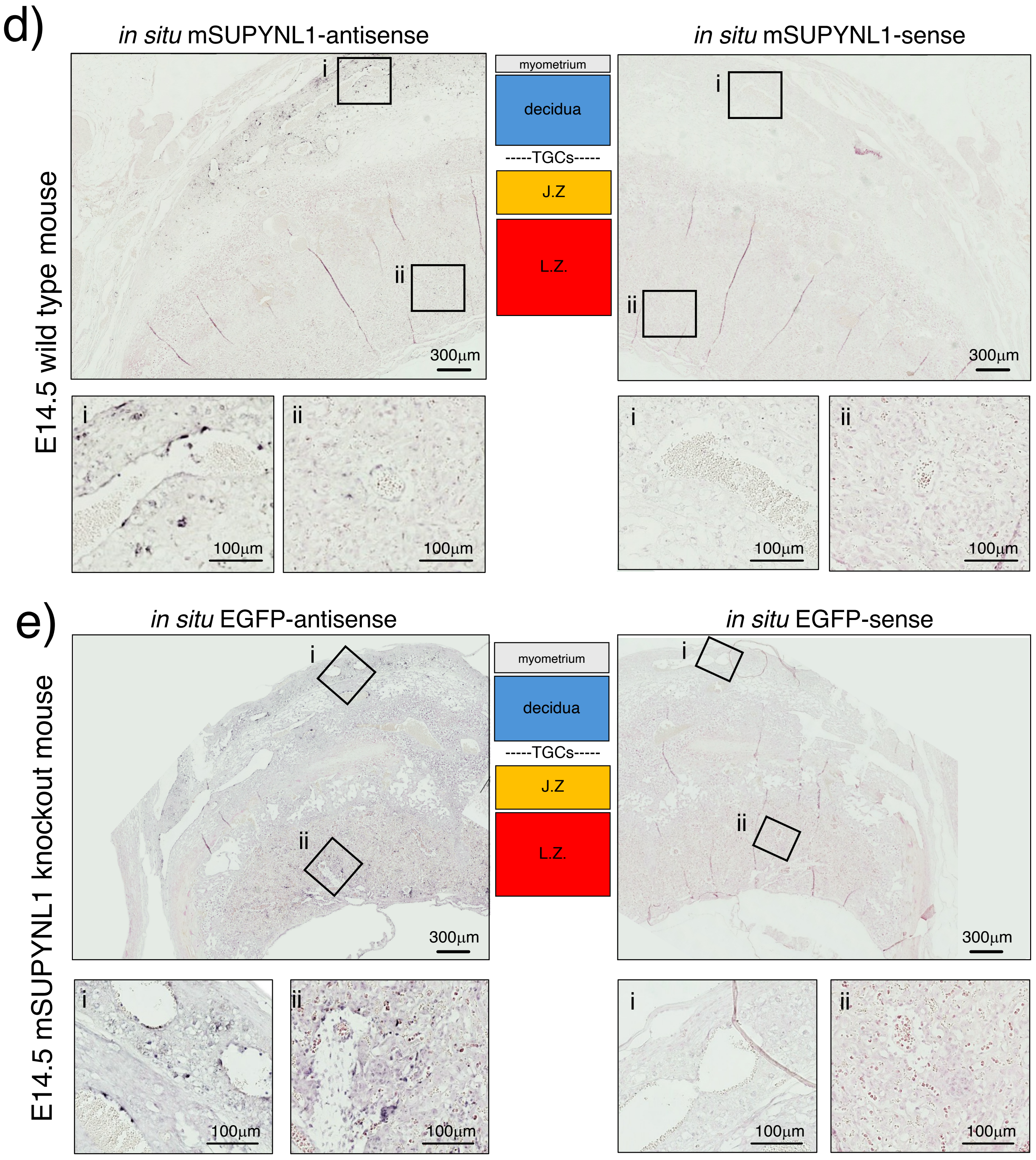


**Supplementary Figure 5. Expression analysis of mSUPYNL1 in wild-type and mSUPYNL1 knockout mice.**

a. Northern blot analysis of mSUPYNL1 mRNA expression in mouse tissues. A commercial mouse multiple-tissue Northern blot membrane (MR-10; GenoStaff, Tokyo, Japan) was hybridized with an mSUPYNL1-specific probe. After detection of mSUPYNL1 transcripts, the membrane was reprobed with an β-actin probe as an internal control. Arrowheads indicate the position of an approximately 3-kb transcript, which is presumed to represent the mSUPYNL1 transcript.

b. RT-PCR analysis of mSUPYNL1 expression in various tissues from wild-type mice and EGFP expression in corresponding tissues from mSUPYNL1 knockout mice. GAPDH was used as an internal control. RT(−) indicates a negative control prepared without reverse transcriptase to exclude the possibility of genomic DNA contamination.

c. In situ hybridization analysis of mSUPYNL1 expression at embryonic day E8.5. Embryos from wild-type mice (left) and mSUPYNL1 knockout mice (right) were hybridized with an mSUPYNL1 antisense probe (left) or an EGFPantisense probe (right), respectively. Higher-magnification images of the boxed regions are shown in the lower panels.

d, e. In situ hybridization analysis of placental and uterine tissues at E14.5. d, wild-type mouse tissues; e, mSUPYNL1knockout mouse tissues. The left panels show hybridization with antisense probes (mSUPYNL1 in wild-type tissues and EGFP in knockout tissues), whereas the right panels show the corresponding sense-probe controls. Higher-magnification images shown below each whole-tissue section correspond to (i) the decidua and myometrium and (ii) the placental labyrinth zone.

| Cloning | | | |
|---|---|---|---|
| gene | primer ID | Forward (5'-3') | Reverse (5'-3') |
| mSUPYNL1 | C11-1st | CCCAGCCCTAGCCCTATAA | CTCGAACTCAGAAATCCGTCTG |
| | C11-2nd(EcoRV-Bcl) | CGGATATCATGAAAGGTGATGAAGTCTC | CCTGATCATAATACGTGTCGGATGGTGGAG |
| mSUPYNL1-SU | C11SUdelTM | CCAGCTGCGGATCCGGTACCAGATTA | CGGATCCGCAGCTGGGAATGGAGAT |
| mSUPYNL1-TM | C11delSUTM | ATATCATGGCTACTGTTGCCGAGATC | CAGTAGCCATGATATCAAGCTTATC |
| syncytin-A | synA-1st | AAGCCTTGCCTTTGGAACTCTCAG | AACAGACTGGGAAGCGAAGTTGGA |
| | synA-2nd(Kpn-Not) | CGGGTACCATGGTTCGTCCTTGGGTTT | TCGCGGCCGCCTAAGCGTAATCTGGAACATCGTAT GGGTAATAGACGGCATCCTCCTCCG |
| syncytin-A SU | synASU-2nd | CGGGTACCATGGTTCGTCCTTGGGTTT | TCGCGGCCGCCTAAGCGTAATCTGGAACATCGTAT GGGTAAGAAGCGGGGACAGAGGAGG |
| syncytin-B | synB-1st | AAGGAATCTCTCACTGGCTGCACT | AACAACTCCAGAGGCCATGTCTGA |
| | synB-2nd(Kpn-Not) | CGGGTACCATGACAGGCTTTTGGGTCC | TCGCGGCCGCCTAAGCGTAATCTGGAACATCGTAT GGGTAATATGTAGGAATGGTGTC |
| syncytin-B SU | synBSU-2nd | CGGGTACCATGACAGGCTTTTGGGTCC | TCGCGGCCGCCTAAGCGTAATCTGGAACATCGTAT GGGTAAAGAACTGAGTCTGAGGAAA |
| syncytin-1 | syn1-1st | AACTGCGGTTAAAGTGGCTGGAGT | TTGGTCAGGTGTGAGCTAAGTTGC |
| | syn1-2nd(EcoRV-Not) | CGGATATCAGGATTTGCGCCTGCTCTTCAAAC | TCGCGGCCGCCTAAGCGTAATCTGGAACATCGTAT GGGTAATAACTGCTTCCTGCTGA |
| syncytin-1 SU | syn1SU-2nd | CGGATATCAGGATTTGCGCCTGCTCTTCAAAC | TCGCGGCCGCCTAAGCGTAATCTGGAACATCGTAT GGGTAGGGCTTAGATATGACATAAC |
| syncytin-2 | syn2-1st | ACTTGTACACCACCAGGAGTTCCA | AGCGGGTGACTTGAGAGATCCAAT |
| | syn2-2nd(Kpn-Not) | CGGGTACCATGGGCCTGCTCCTGCTGG | TCGCGGCCGCCTAAGCGTAATCTGGAACATCGTAT GGGTAATAGAAGGGTGACTCTTG |
| syncytin-2 SU | syn2SU-2nd | CGGGTACCATGGGCCTGCTCCTGCTGG | TCGCGGCCGCCTAAGCGTAATCTGGAACATCGTAT GGGTAGGGCAACGGGGAATTCCCAT |

| RT-PCR | | | |
|---|---|---|---|
| gene | primer ID | Forward (5'-3') | Reverse (5'-3') |
| mSUPYNL1 | C11-RT- | AGGGTCTCCTCAGAGTGATT | AGCATCTCCCGTCTAGTGAT |
| syncytin-A | synA-RT- | GAGTTGAGGCAGAAGGAGTATG | CTGGGAATATGAACCCACTGTTA |
| syncytin-B | synB-RT- | GAACGAGATCACTGTCCAAGAA | CCTGAAGAGTTGAGGCAGAAG |
| EGFP | EGFP-RT- | ACGTAAACGGCCACAAGTTC | TGCTCAGGTAGTGGTTGTCG |
| mGAPDH | GAPDH-RT- | GCATCCTGGGCTACACTGAG | TCCACCACCCTGTTGCTGTA |

| Knock out mouse | | | |
|---|---|---|---|
| gene | primer ID | Forward (5'-3') | Reverse (5'-3') |
| CRISPER target-left | target(L) | TGAAGTCTCGATCACTAGAC | |
| CRISPER target-right | target(R) | CGTATTAAAATGCTAACCCT | |
| Knock-In donor-5' | KI-5arm | CGGTACCCGGGGATCGAAAACGAAAGCCCTAGTGT | GCCCTTGCTCACCATAGTGATCGAGACTTCATCAC |
| Knock-In donor-EGFP | KI-EGFP | GAAGTCTCGATCACTATGGTGAGCAAGGGC | CGACTCTAGAGGATCCTGCAGGTCGAGGGA |
| Knock-In donor-3' | KI-3arm | GCAGGCATGCAAGCTCCTTGGATTCTGCTCAAGGG | TGATTACGCCAAGCTTCTTAGGCTGTGTGGGTGAT |

**Supplementary Table 1. Primer sequences used in this study.**

| | Species | Clone No. | Cat No. | Distributor | Dilution | | |
|---|---|---|---|---|---|---|---|
| | | | | | IHC | ICC | WB |
| mouse suppressyn-like1 | Rabbit | C11b | n.a. | n.a. | | | 1/2500 |
| ASCT2 | Rabbit | D7C12 | 8057 | CST | | | 1/2000 |
| Anti-DDDDK-tag mAb | Mouse | FLA-1 | M185 | MBL | | 1/1000 | 1/10000 |
| Anti-DDDDK-tag pAb | Rabbit | n.a. | PM020 | MBL | | | 1/3000 |
| β-actin | Mouse | AC-15 | A5441 | Sigma | | | 1/10000 |
| Anti-HA-tag mAb | Mouse | TANA2 | M180-3 | MBL | | | 1/5000 |
| Anti-HA-tag pAb | Rabbit | n.a. | 561 | MBL | | | 1/3000 |
| Anti-GFP(Green Fliorescent Protein)pAb | Rabbit | n.a. | 598 | MBL | | | 1/2500 |
| Anti-HTLV1-gp46 | Mouse | 1C11 | sc-53890 | SantaCruz | | | 1/2000 |
| VimentinXP Rabbit mAb | Rabbit | D21H3 | 5741 | CST | 1/1000 | | |
| Perforin Polyclonal antibody | Rabbit | n.a. | 14580-1-AP | proteintech | 1/5000 | | |
| Anti-CD31 antibody | Rabbit | n.a. | ab124432 | abcam | 1/3000 | | |
| DBA-Biotin | n.a. | n.a. | J204 | MGC | 1/1000 | | |
| Alexa Fluor 488 goat anti-mouse IgG(H+L) | Mouse | n.a. | A11029 | Thermo Scientific | | 1/1000 | |
| Anti-mouse IgG HRP-linked Antibody | Mouse | n.a. | 7076 | CST | | | 1/5000 |
| Anti-rabbit IgG HRP-linked Antibody | Rabbit | n.a. | 7074 | CST | | | 1/5000 |
| Mouse TrueBlot ULTRA Anti-Mouse Ig HRP | Mouse | n.a. | 18-8817-30 | ROCKLAND | | | 1/2500 |

**Supplementary Table 2. Antibody list used in this study.**

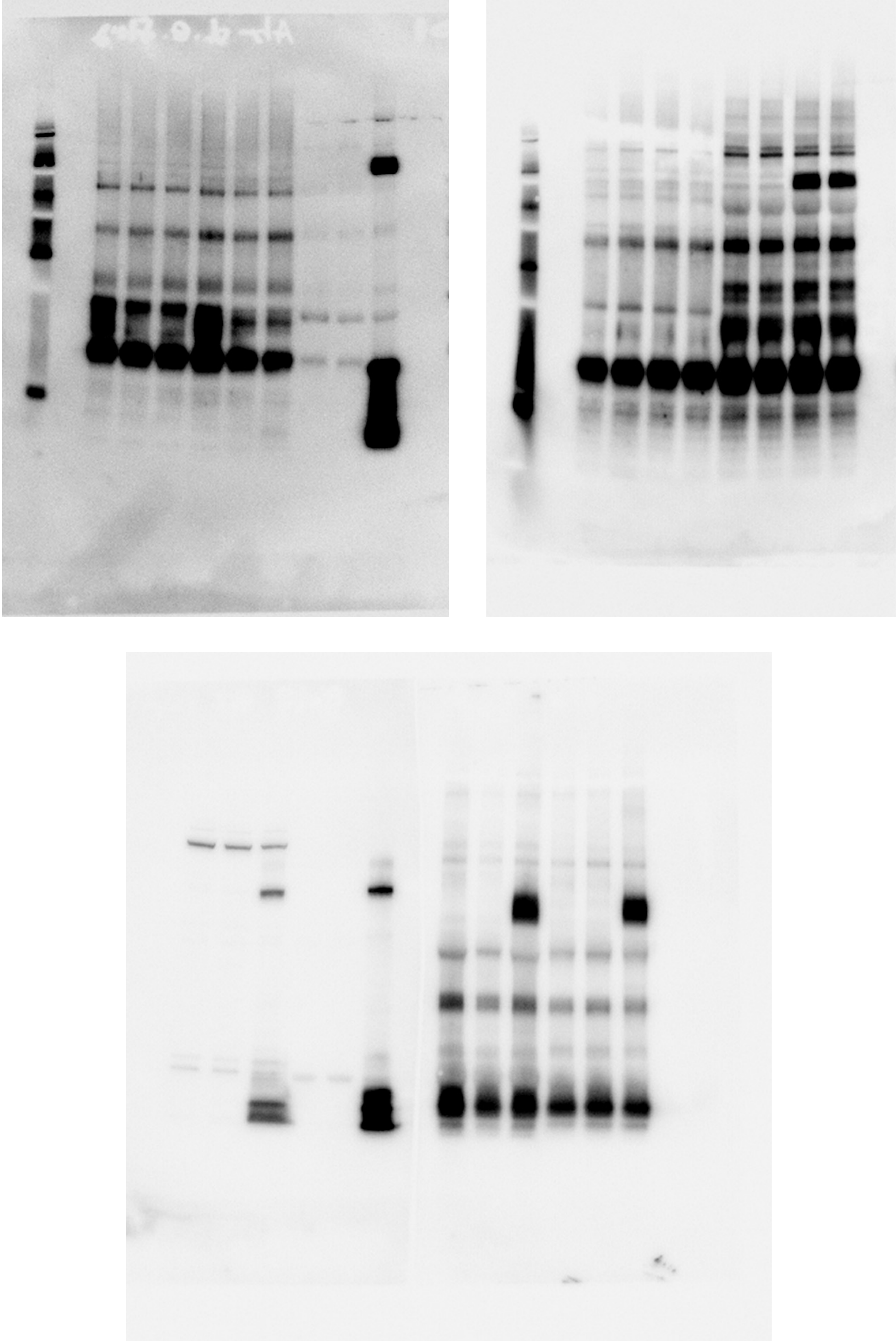

**Supplementary Figure 6. Uncropped original immunoblot images used in Figure 1b.**

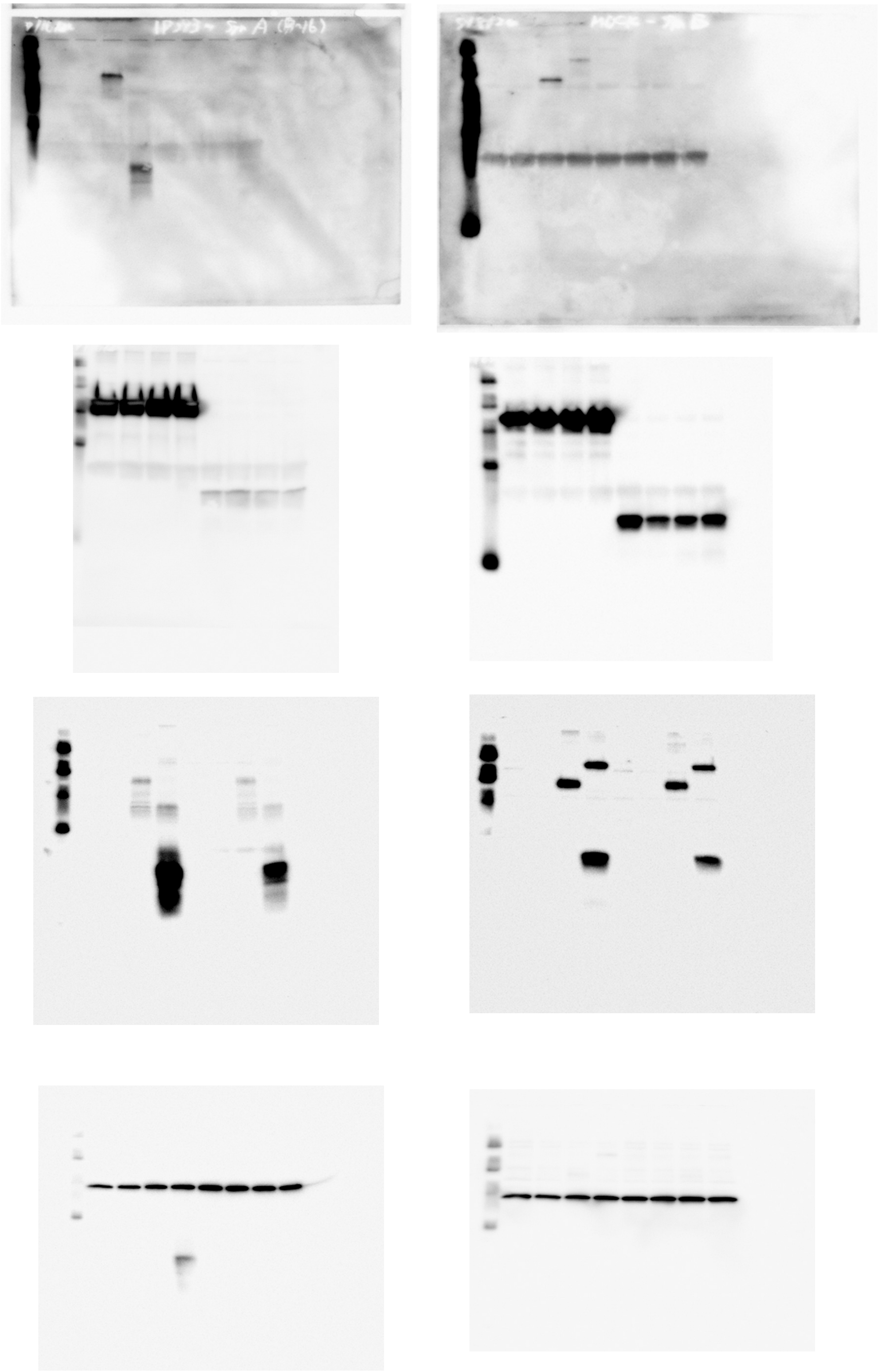

**Supplementary Figure 7. Uncropped original immunoblot images used in Figure 2c.**

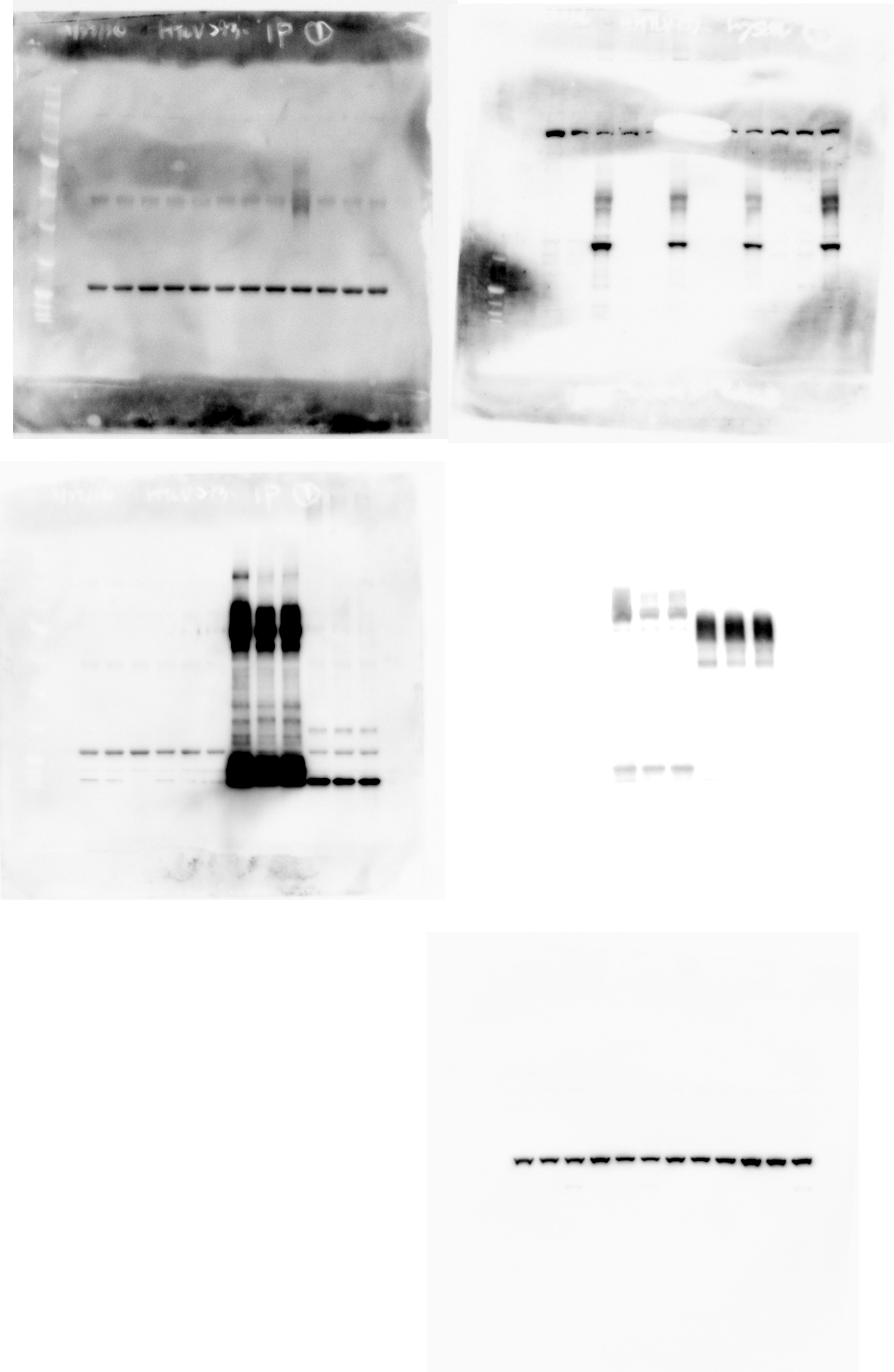

**Supplementary Figure 8 Uncropped original immunoblot images used in Figure 4d.**

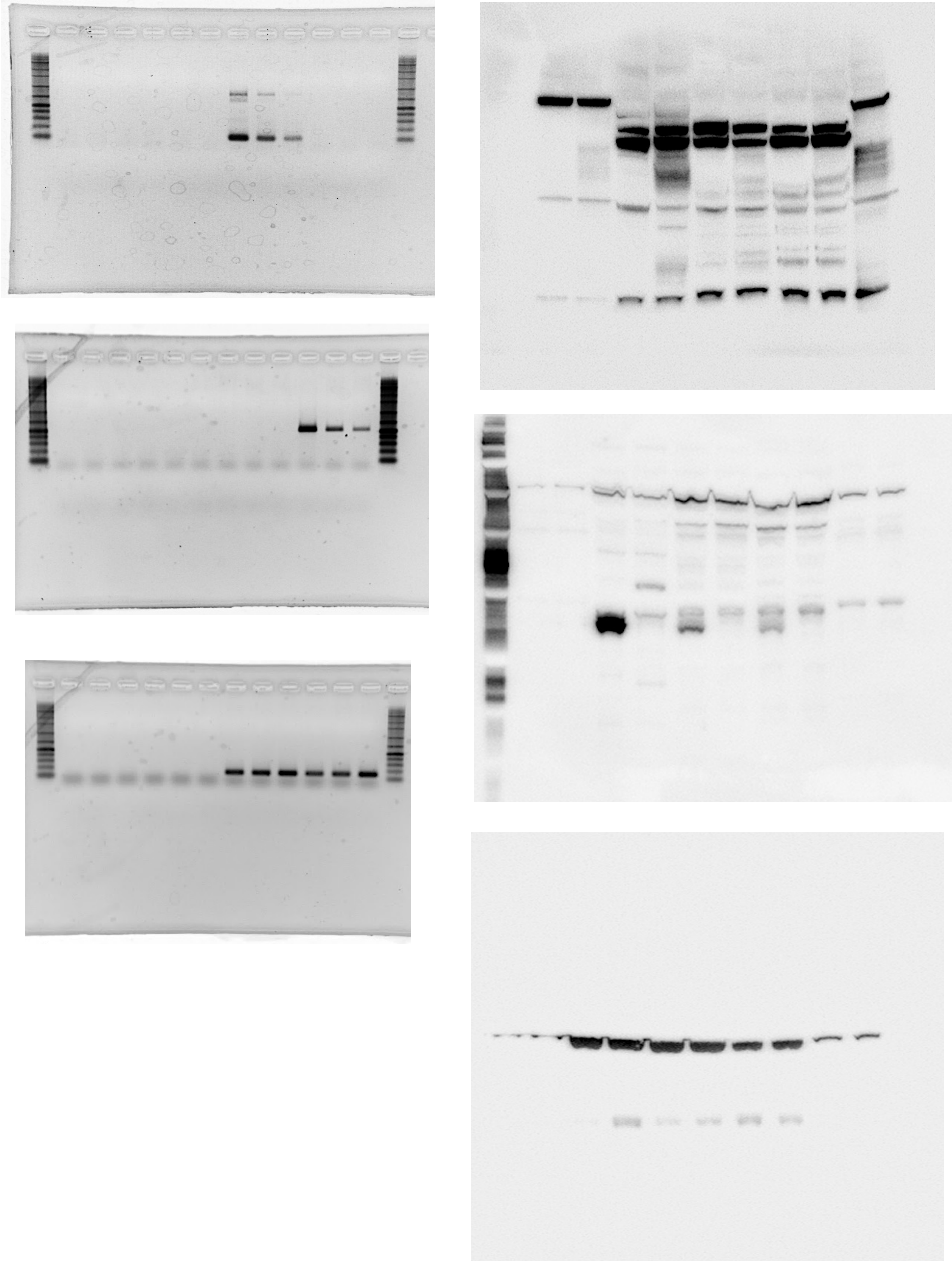

**Supplementary Figure 9. Uncropped original Gel image or immunoblot images used in Figure 3b,c.**
